# Effective purifying selection in ancient asexual oribatid mites

**DOI:** 10.1101/112458

**Authors:** Alexander Brandt, Ina Schaefer, Julien Glanz, Tanja Schwander, Mark Maraun, Stefan Scheu, Jens Bast

**Author notes:** Corresponding authors: Alexander Brandt, Georg-August-University, J.-F.-Blumenbach Institute of Zoology and Anthropology, Berliner Str. 28, DE - 37073 Göttingen; Jens Bast, University of Lausanne, Department of Ecology and Evolution, UNIL Sorge, Le Biophore, CH - 1015 Lausanne.

## Abstract

Sex is beneficial in the long-term, because it can prevent mutational meltdown through increased effectiveness of selection. This idea is supported by empirical evidence of deleterious mutation accumulation in species with a recent transition to asexuality. Here, we studied the effectiveness of purifying selection in oribatid mites, which have lost sex millions of years ago and diversified into different families and species while reproducing asexually. We compared the accumulation of deleterious coding and non-coding mutations between three asexual and three sexual lineages using transcriptome data. Contrasting studies of young asexual lineages, we find evidence for strong purifying selection that is more effective in asexual compared to sexual oribatid mite lineages. Our results suggest that large populations likely sustain effective purifying selection and facilitate the escape of mutational meltdown in the absence of sex. Thus, sex *per se* is not a prerequisite for the long-term persistence of animal lineages.

---

One of the most challenging problems in evolutionary biology is to explain the maintenance of sex ^1,2^. Albeit sex is coupled with substantial costs, including the production of males, transmission disadvantages and costs associated with mating and recombination, the mode of reproduction for the great majority of animal species is obligate sex ^3–5^. There is little agreement on which mechanisms generate selection for sex in natural populations in the short-term ^6,7^. However, it is established consensus that sex and recombination are beneficial for the long-term persistence of lineages as based on theoretical predictions and empirical evidence for reduced purifying selection in asexual eukaryotes ^8,9^.

Comparing rates of deleterious mutation accumulation in natural populations of sexual and asexual organisms is difficult as other factors than sex and recombination affect the effectiveness of selection. Most importantly, it is also determined by population size and mutation rate ^10,11^. Nevertheless, empirical estimates showed increased accumulation of deleterious coding mutations in effectively clonal lineages (hereafter referred to as asexual) as compared to closely related sexuals, e.g. in natural populations of *Timema* stick insects, *Campeloma* and *Potamopyrgus* freshwater snails and *Oenothera* evening primroses ^12–16^. Coding mutation accumulation occurred at up to 13.4 times the rate in sexuals. Further, there is evidence for accumulation of coding and non-coding mutations in non-recombining genomic regions of sexual organisms, such as mitochondria or Y-chromosomes ^17,18^.

All asexual lineages that were analysed for mutation accumulation lost sex relatively recently between 5,000 years and 1.5 my ago. Whether similar patterns are observed in long-term asexual populations, i.e. after tens of millions of years without sex remains an open question. Few asexual lineages are known that have persisted and diversified in the absence of sex, most prominently bdelloid rotifers, darwinulid ostracods and various clades of oribatid mites ^19–22^. Two studies investigated the accumulation of deleterious coding mutations in bdelloid rotifers ^23,24^, but recent investigations indicate that bdelloid rotifers engage in non-canonical forms of sex and cannot be considered as effectively clonal ^25–27^. Further, there are no studies on mutation accumulation for darwinulid ostracods, only evidence of very slow background mutation rates ^28^. Oribatid mites lost sex multiple times independently, several million years ago, followed by extensive radiation of parthenogenetic clades as indicated by their phylogenetic distribution and high inter- and intraspecific divergence ^29–33^. Thus this speciose, largely soil-living animal group (~10,000 species ^19^) is well suited for comparative investigations of genomic consequences under long-term asexuality.

Here, we tested if long-term asexuality in oribatid mites resulted in signatures of reduced effectiveness of selection by comparing the accumulation of deleterious coding mutations and changes in codon usage bias. We analysed nuclear and mitochondrial orthologous genes from newly generated transcriptomic data of the three sexual species *Achipteria coleoptrata* (Linnaeus, 1758), *Hermannia gibba* (Koch, 1839) and *Steganacarus magnus* (Nicolet, 1855) and the three parthenogenetic species *Hypochthonius rufulus* (Koch, 1835), *Nothrus palustris* (Koch, 1839) and *Platynothrus peltifer* (Koch, 1839). Additionally, we analysed two nuclear genes (*hsp82* and *ef1α*) of 30 species (9 sexual, 21 asexual species) from different phylogenetic groups ^34^ (Fig. 1). We found no evidence for reduced purifying selection in asexual lineages, neither for elevated accumulation of deleterious coding point mutations nor for relaxed selection on codon usage bias. Surprisingly, our results indicate even more effective purifying selection in asexual as compared to sexual oribatid mites.

**Figure 1:**
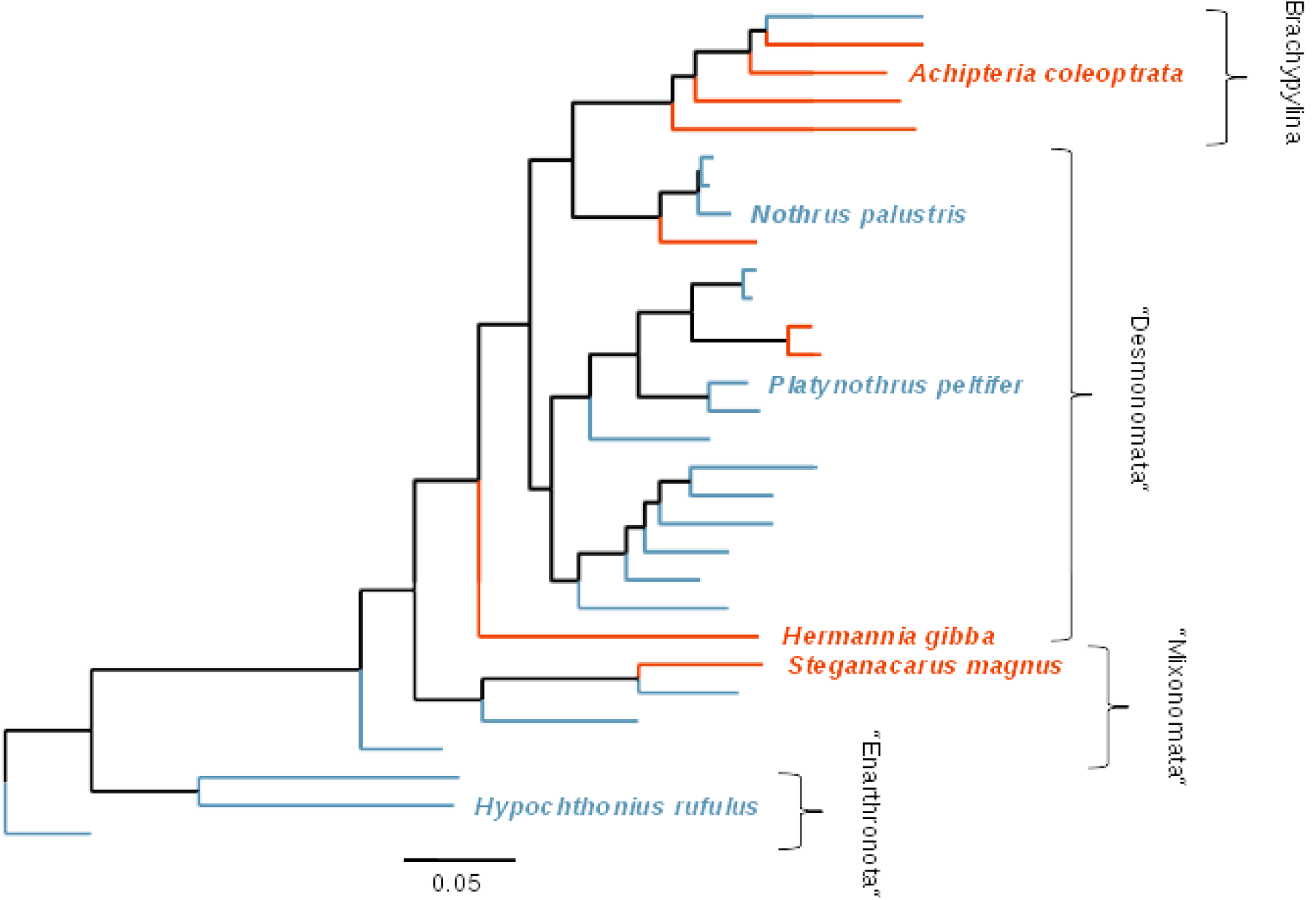
Phylogenetic tree based on 18S rDNA, *ef1α* and *hsp82* of 30 oribatid mite species with asexual (blue) and sexual (red) species analysed in this study (see Methods). The taxon sampling covers species from four out of six major groups of oribatid mites (brackets). Species used for transcriptome-wide analyses are indicated with full name, remaining species names are listed from top to bottom in Supplementary Table 1.

## Results

We used two approaches to compare the effectiveness of purifying selection between asexual and sexual oribatid mite lineages on the transcriptome scale. First, we analysed the accumulation of coding point mutations and inferred their potential ‘deleteriousness’. Second, we inferred the effectiveness of selection on codon usage bias since purifying selection also acts on synonymous sites ^35^. Analyses are based on 3,545 nuclear orthologous genes shared among the six species (see Methods for details).

## Asexuals accumulate coding mutations more slowly than sexuals

To estimate the accumulation rate of coding point mutations in orthologs, we computed the ratio of non-synonymous (coding) to synonymous (non-coding) divergence (dN/dS) as a measure of amino acid changes normalised for background mutation rates (see Methods). Given the long divergence time between the six species used for sequencing of transcriptomes (> 200 myr ^36^), detectable orthologs shared among the six species are expected to be under strong purifying selection (dN/dS < 1). In genes under purifying selection any coding change is likely deleterious, hence an increased accumulation of deleterious coding mutations results in a higher dN/dS ratio ^37^. We analysed coding mutation accumulation exclusively at terminal branches of the phylogenetic tree because character states (i. e. sexual or asexual reproduction) at internal branches are uncertain for methodological reasons ^38^. We used a model of codon evolution estimating one dN/dS ratio for internal, one for terminal branches of asexual species and one for terminal branches of sexual species (from here referred to as terminal asexual and sexual branches, respectively; three-ratio model; see Methods). As expected, all 3,545 orthologs were under purifying selection at terminal branches (mean dN/dS = 0.084). Surprisingly and contrary to expectations, per-gene dN/dS ratios were on average lower at asexual branches as compared to sexual branches (mean Δ_asex-sex_ = -0.008; Wilcoxon signed-rank test *P <* 0.001; Fig. 2). Non-coding substitution rates (dS) did not differ between reproductive modes (means of 1.112 and 1.146 for asexual and sexual branches, respectively; gene effect *P* = 0.951, reproductive mode effect *P* = 0.218, interaction *P* = 0.999; permutation ANOVA), indicating that the differences in dN/dS ratios between reproductive modes were not driven by differences in dS.

**Figure 2:**
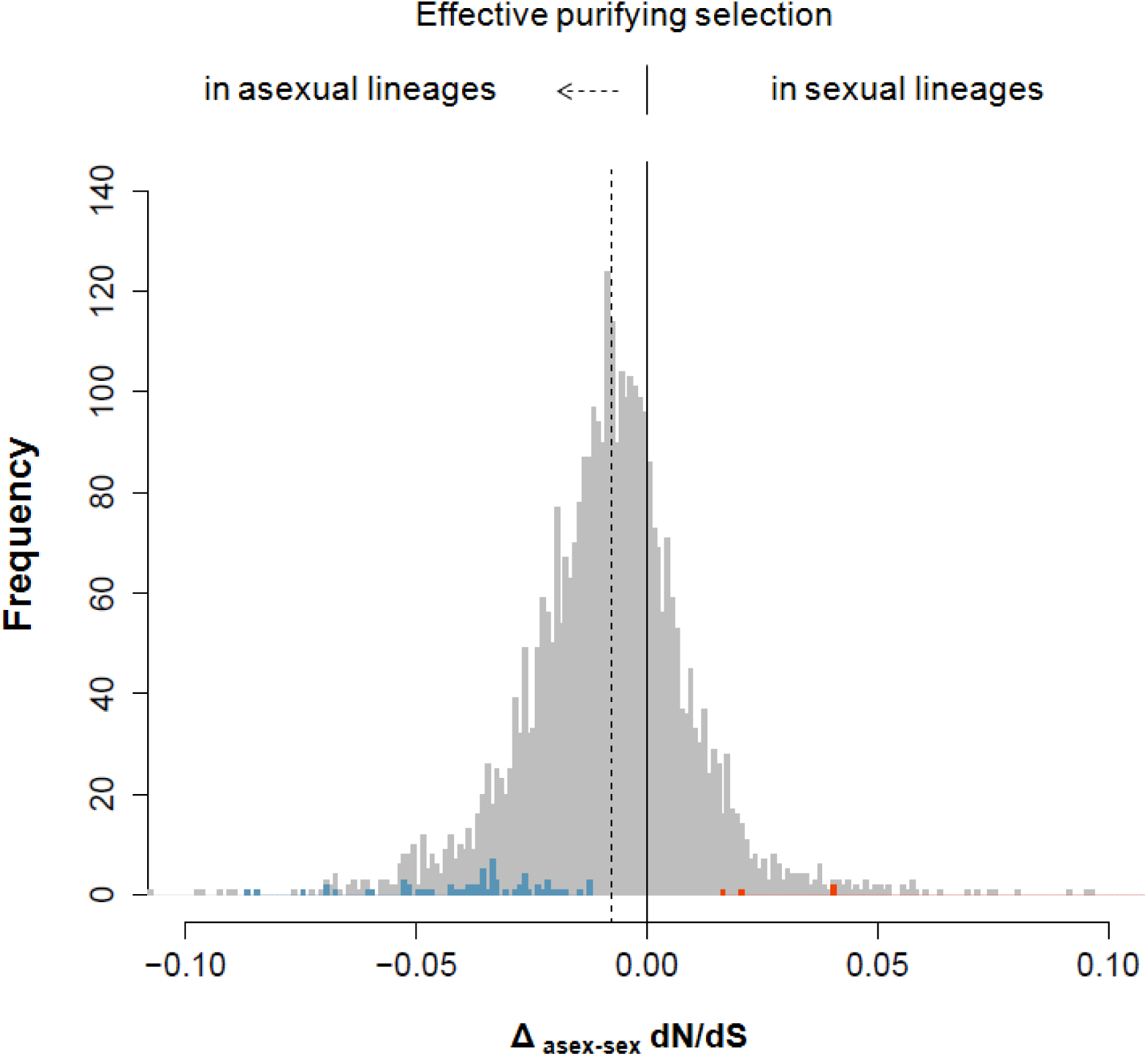
Histogram showing the frequencies of per-gene differences in dN/dS ratios between asexual and sexual terminal branches in a tree comprising three sexual and three asexual oribatid mite species for 3,545 orthologous loci under purifying selection (dN/dS < 1). The histogram range is limited between -0.1 and 0.1. The distribution mean (-0.008; dotted line) is shifted to the left indicating overall lower dN/dS ratios at asexual as compared to sexual branches (Wilcoxon signed-rank test *P* < 0.001). Frequencies of genes with significantly lower dN/dS ratios at asexual as compared to sexual branches or sexual as compared to asexual branches are colored blue and red, respectively.

Although orthologs were overall under stronger purifying selection in asexual compared to sexual oribatid mites, dN/dS values as well as their difference between asexuals and sexuals varied widely among orthologs (ranges: dN/dS 0-0.627; Δ_dN/dS_ 0-0.540). We therefore also analysed which specific orthologs differed significantly in their effectiveness of purifying selection between reproductive modes.

To identify orthologs under significantly stronger purifying selection in asexual and sexual lineages, we tested if the three-ratio model was a better fit to the data than a model allowing for one dN/dS ratio for terminal and internal branches, without discriminating between reproductive modes (two-ratio model; see Methods). For 67 orthologs dN/dS ratios were significantly lower at asexual as compared to sexual branches and for five orthologs at sexual as compared to asexual branches (colored bars; Fig. 2). The orthologs with strong selection in asexual mites were enriched for Gene Ontology terms related to e.g. immune response, mitotic cell cycle and germline cell division (Supplementary Data 1).

## Coding mutations are less deleterious in asexuals as compared to sexuals

In addition to elevated rates of coding mutation accumulation, reduced purifying selection is expected to translate into coding mutations with strong deleterious effects.

We therefore analysed the potential deleteriousness of the amino acid substitutions. For this, we inferred changes in hydrophobicity scores (HS) between ancestral and replacement amino acids as a measurement of the “deleteriousness” of coding mutations (see Methods) because protein folding is influenced by changes in hydrophobic properties at amino acid substitution sites ^39^. For nuclear orthologs, transitions between amino acids with more dissimilar hydrophobicity (HS < 90) involved 46,857 changes (50.74% of overall changes) at asexual and 65,491 (50.98%) at sexual branches. In accordance with the faster coding mutation accumulation in sexual than asexual oribatid mites reported above, amino acids shifted to more dissimilar hydrophobicity at sexual branches (glmm *z* = 2.4, *P* = 0.0171; Fig 3a), indicating stronger deleteriousness of coding changes in sexual as compared to asexual oribatid mites.

**Figure 3:**
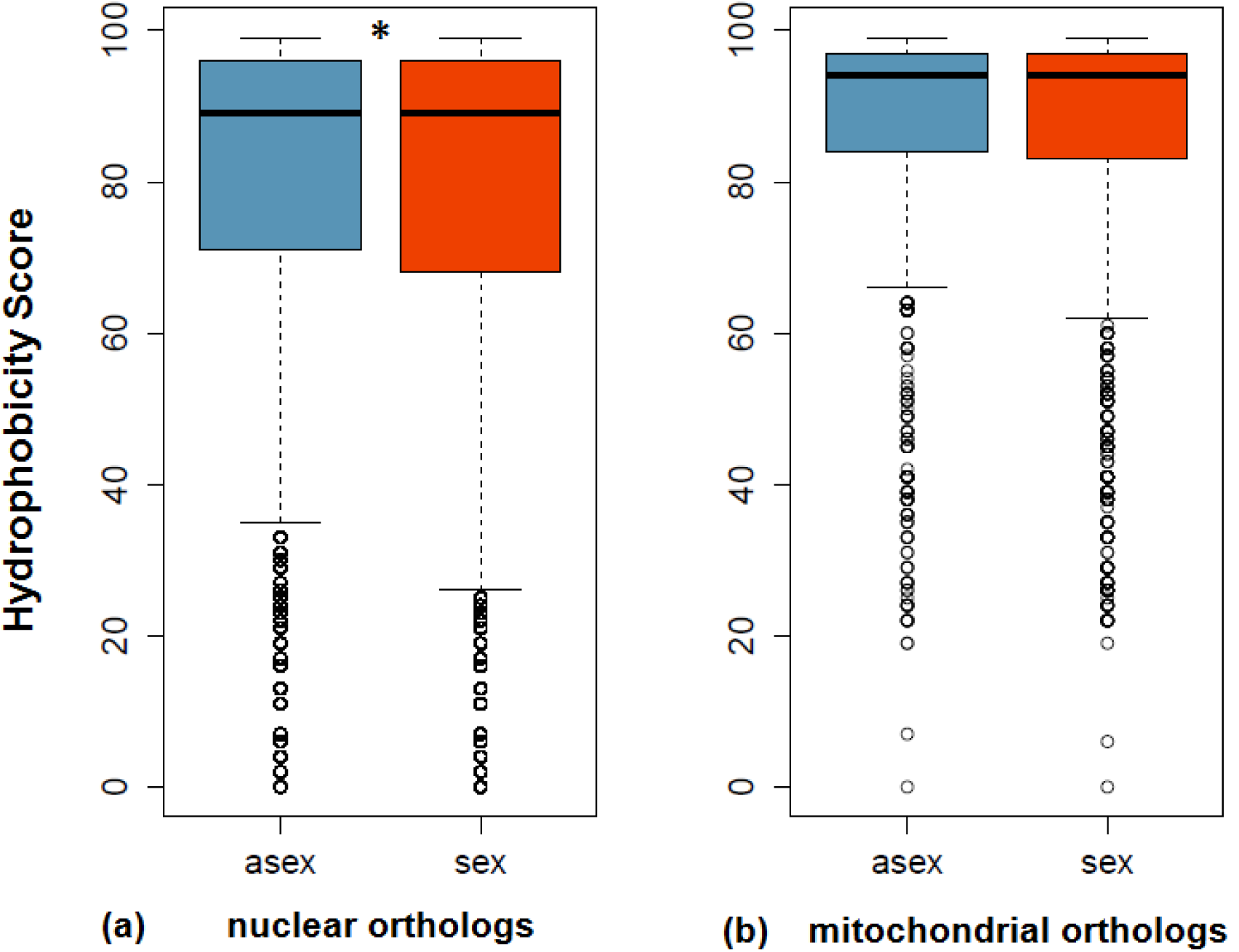
Boxplots of Hydrophobicity Scores (HS) at three asexual (blue) and three sexual (red) terminal branches for (a) 3,545 nuclear and (b) 10 mitochondrial orthologous genes. HS measures the strength in changes of hydrophobicity from ancestral to replacement substitutions and indicates the “deleteriousness” of a coding mutation. The lower the HS the stronger the change in hydrophobicity and “deleteriousness” of a coding mutation. Significant differences in HS are marked with an asterisk (GLMM; \**P* <0.05; see Methods).

## Results on the transcriptome scale are supported by a larger taxon sampling

Overall, based on the analyses of transcriptomic orthologs shared among six species, asexual oribatid mite lineages accumulate less deleterious coding mutations as compared to sexual lineages. However, the small number of taxa used can generate branch length uncertainties which could affect inferences of dN/dS and ancestral amino acid sequences. To reduce this problem, we extended the taxon sampling to a total of 30 species. Using the same approaches, we compared coding mutation accumulation among 21 asexual and 9 sexual oribatid mite species (Fig. 1, Supplementary Table 1) in two nuclear genes *(ef1α* and *hsp82*) generated in a previous study ^34^. Both genes were under strong purifying selection at terminal branches as indicated by dN/dS ratios (0.045 and 0.034 for *ef1α* and *hsp82*, respectively). There were no significant differences between asexual and sexual branches as the two-ratio model provided a better fit to the data than the three-ratio model (Table 1). Non-coding substitution rates (dS) did not differ between reproductive modes, (*P* = 0.540 and *P* = 0.237 for *ef1α* and *hsp82*, respectively). This lack of a significant difference between dN/dS ratios is unlikely the result of low variation in *ef1α* and *hsp82*, as mean pairwise divergence of amino acid sequences was high in both genes (0.069 and 0.109 for *ef1α* and *hsp82*, respectively). Consistent with dN/dS ratios, there were no differences in hydrophobicity changes at asexual and sexual branches for both genes (*P* = 0.542 and *P* =0.624 for *ef1α* and *hsp82*, respectively; Wilcoxon Rank-Sum test).

**Table 1:**
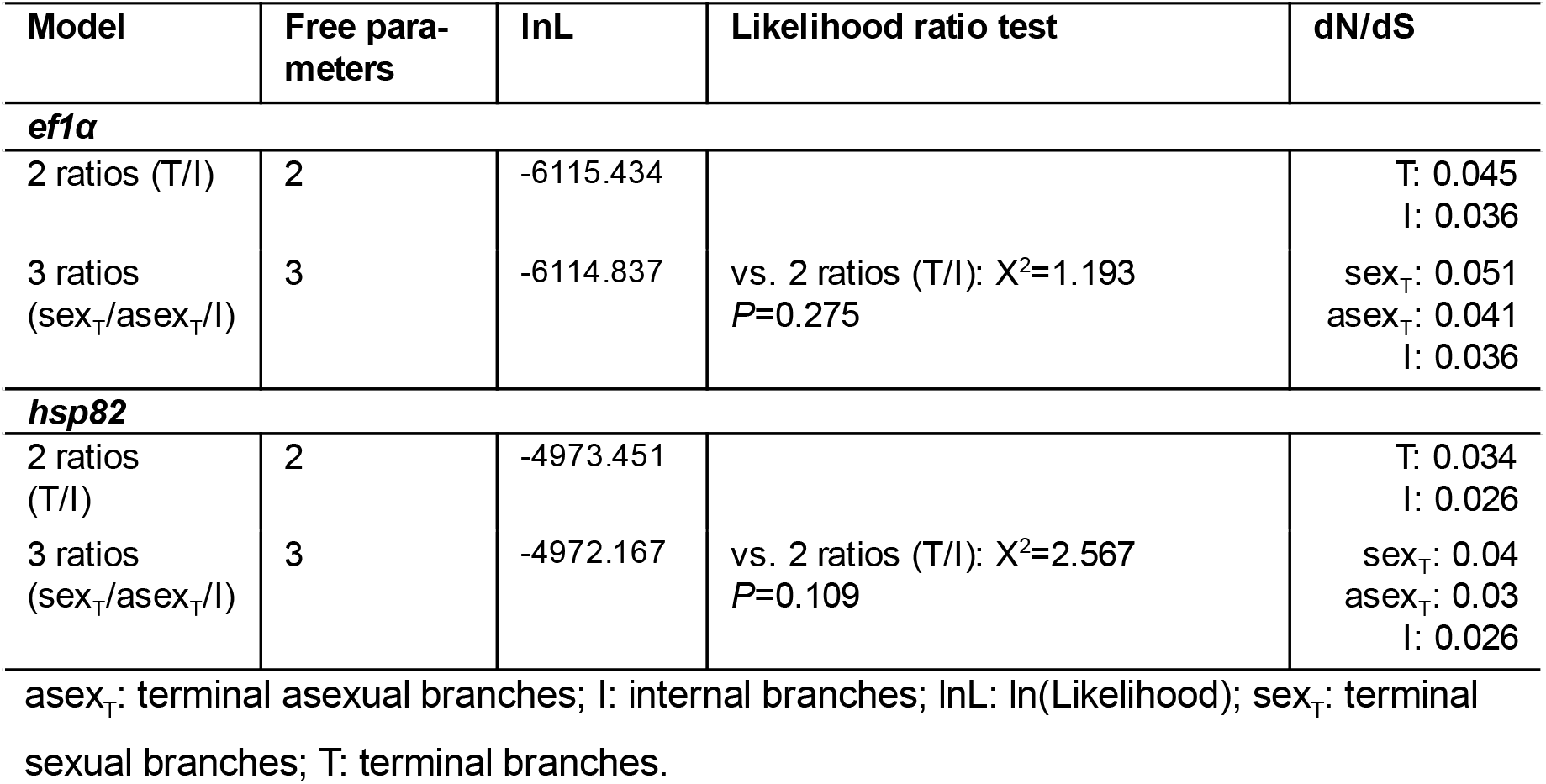
Results of the branch-specific dN/dS ratio analyses of *ef1α* and *hsp82* for 21 asexual and 9 sexual oribatid mite species. Non-significant *P*-values indicate that the 2-ratio model provides the better fit to the data than the 3-ratio model for both genes. Therefore asexual and sexual branches do not differ in these two genes in the effectiveness of selection on coding sites.

As indicated by the transcriptome analyses described above, the dN/dS ratio differences between sexual and asexual taxa varied considerably among orthologs and only analyses of large gene sets are likely to reveal putative differences between reproductive modes. Indeed the gene *ef1α*, although present in the transcriptome ortholog set, did not feature significantly different dN/dS ratios between reproductive modes (0.015 and 0.026 at asexual and sexual branches, respectively; *P* = 0.076). *Hsp82* was absent from the ortholog dataset, preventing estimation of dN/dS ratios from the transcriptomes for this gene.

### Asexuals accumulate non-coding mutations more slowly than sexuals

Although mutations at non-coding sites are generally considered to be neutral, they can be under selection for example because different codons influence the speed and accuracy of protein translation ^35^. Hence, we also assessed the effectiveness of selection acting at non-coding sites by estimating codon usage bias (CUB). This method is particularly robust, because measurements of CUB do not depend on likelihood estimates, tree topologies or branch length estimates. As the metric, we used the codon deviation coefficient (CDC ^40^), which calculates the deviation from expected CUB and accounts for background nucleotide composition, thus allowing for cross-species comparisons. A lower CDC value indicates more ‘relaxed’ selection on CUB (see Methods). We compared gene-specific CDC values for each of the 3,545 orthologous nuclear genes between reproductive modes. Consistent with the results for nuclear dN/dS ratios, per-gene CDC was slightly but significantly higher in asexual species (Fig. 4a; means of 0.130 and 0.128 for asexual and sexual species, respectively; gene effect *P* < 0.001, reproductive mode effect *P* = 0.008, interaction *P* = 0.616; permutation ANOVA), indicating more effective purifying selection at non-coding sites in asexuals.

**Figure 4:**
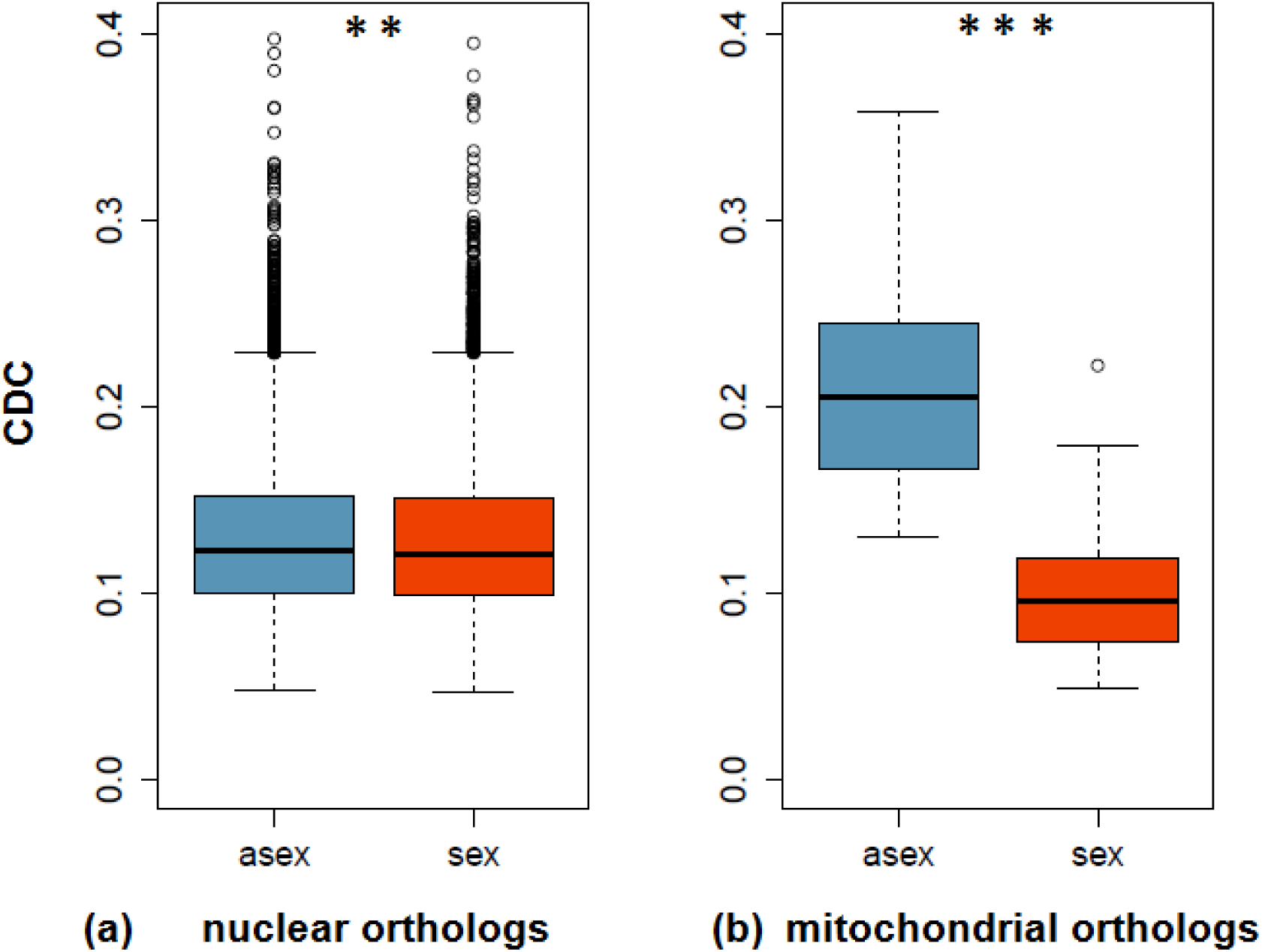
Boxplots of CDC values of three asexual (blue) and three sexual (red) oribatid mite species for (a) 3,545 nuclear and (b) 10 mitochondrial orthologous genes. CDC measures the deviation of observed from expected codon usage and allows for inference of the effectiveness of purifying selection acting on non-coding sites. A lower CDC value corresponds to more ‘relaxed’ selection on codon usage bias (see Material and Methods). Significant differences in CDC are marked with asterisks (as inferred with permutation ANOVAs (n=5000); ^***^ *P* <0.001 and ^**^ *P* <0.01; see Methods).

### More effective selection in asexuals for non-coding mitochondrial sites

Next, we analysed mitochondrial genes, which allows for comparing the effectiveness of purifying selection among non-recombining genomes with different nuclear genomic backgrounds. We analysed the effectiveness of purifying selection in 10 orthologous mitochondrial genes (*atp6, cob, cox1, cox2, cox3, nd1, nd2, nd3, nd4, nd5*) using the same approaches as for nuclear genes. Surprisingly, and contrasting to the results of nuclear orthologs, dN/dS ratios were lower at terminal sexual branches as compared to terminal asexual branches (mean Δ_asex-sex_ = 0.027; Wilcoxon signed-rank test *P* = 0.006). However, dS rates were highly elevated for sexual as compared to asexual branches (means of 2.814 and 5.577 for asexual and sexual branches, respectively; gene effect *P* = 0.044, reproductive mode effect *P* = 0.002, interaction *P* = 0.876; permutation ANOVA). This raises the question, if the observed differences in mitochondrial dS between reproductive modes could be influenced by selection at non-coding sites ^41^. Indeed, non-coding mitochondrial substitution rates (dS) and selection at non-coding sites as measured by CDC were negatively correlated in sexual lineages (r = -0.37, *P* = 0.042) but not in asexual lineages (r = 0.16, *P* = 0.396, Spearman’s rank correlation). Because dS rates strongly differed and were adversely influenced by selection at non-coding sites between reproductive modes, mitochondrial dN/dS ratios are not comparable. Based on dN rates only, there was no difference in asexual and sexual coding changes (means of 0.173 and 0.231 and for asexual and sexual branches, respectively; gene effect *P* = 0.037, reproductive mode effect *P* = 0.138, interaction *P* = 0.994; permutation ANOVA). Consistent with dN rates, the analysis of hydrophobicity changes (‘deleteriousness’) of amino acids for mitochondrial orthologs showed no overall difference between reproductive modes (glmm *z* = 0.81, *P* = 0.421; Fig. 3b). At mitochondrial non-coding sites, effectiveness of selection was strongly reduced for mitochondria in sexual lineages as compared to those in asexual lineages (Fig 4b; mean CDC of 0.211 and 0.104 for asexual and sexual species, respectively; gene effect *P* = 0.002, reproductive mode effect *P* < 0.001, interaction *P* = 0.9604; permutation ANOVA).

## Discussion

In metazoans, it is believed that sex is important for the long-term persistence of a lineage, because it facilitates effective purifying selection and rapid adaptation despite the effect of drift in finite populations. In contrast to this established consensus, our study shows no evidence for an accumulation of deleterious coding and non-coding point mutations in ancient asexual oribatid mites. Moreover, our results suggest that purifying selection acts even more effectively in asexual as compared to sexual oribatid mite lineages. It is thus unlikely that asexual oribatid mite lineages drift towards extinction because of mutation accumulation. While our taxon sampling for dN/dS ratio estimates of nuclear orthologs in the transcriptome scale analyses was small, the overall pattern is substantiated by results from the more extended taxon set (Figs. 1, 2, Table 1). Moreover, non-coding substitution analyses (CDC, Fig. 4) are independent of assumptions about phylogenetic relationships among taxa and these analyses likewise support the result of more effective purifying selection in asexuals.

Our finding of more effective purifying selection despite the lack of sex is unique among investigated asexual eukaryotes. Previous studies that compared sexual and asexual eukaryotes typically found increased rates of coding mutation accumulation in asexual as compared to sexual lineages, as predicted by theory (reviewed by Hartfield 2015 ^16^). Where no difference was found, sex was lost probably too recently to detect an accumulation of mutations ^42,43^. Increased rates of coding mutation accumulation and relaxed codon usage bias further extends to self-fertilising eukaryotes, although selfers often maintain low rates of outcrossing ^16,44,45^.

What mechanisms could account for the escape of asexual oribatid mites from the ‘mutational meltdown’? Comparing the peculiarities of oribatid mites with those of animal groups including other old asexual lineages may help resolving this question. Oribatid mites are small animals (150-1400 μm) with a wide geographical distribution ^46^. Both aso holds for other old asexual groups, such as darwinulid ostracods and meloidogyne root-knot nematodes, and contrasts most asexual groups that accumulate deleterious mutations ^47,48^. Generally, small body size and large geographic range are indicative of high abundances of organisms ^49,50^, which suggests that large population sizes might alleviate the negative effects of loss of sex.

Traditional models that predict mutation accumulation in asexuals are based on population sizes that are typical for many macroorganisms, but hence might not be applicable to asexuals with large populations ^8,9,51^. Indeed, large population sizes were proposed to maintain effective purifying selection in the absence of sex ^52,53^, because the speed of mutation accumulation in asexual populations is predicted to drop substantially with increasing population sizes and eventually to become biologically irrelevant ^10,11,54–56^. Consistent with the idea that large population size can protect asexual lineages from mutation accumulation, total oribatid mite densities are often very large and can reach up to ~ 350,000 ind./m^2^ in temperate and boreal forests. At high densities (> 100,000 ind/m^2^) asexual species typically contribute > 55% to total species number but > 80% to total density ^46^, indicating substantially larger population sizes of asexual as compared to sexual oribatid mites. This observation is reflected by our findings of purifying selection being more effective in asexual oribatid mite lineages as compared to sexuals in both the nucleus and mitochondria. The observation of a strong difference for selection at non-coding sites for non-recombining mitochondria in sexual and asexual nuclear backgrounds supports that asexual oribatid mite populations are larger than sexuals (Fig 4b). Remarkably, both animal population sizes and the frequency of parthenogenetic reproduction are generally high in soil animals (including protozoans, nematodes, enchytraeids, collembolans, isopods and oribatid mites ^46,57^), suggesting that more ancient asexual scandals may be hidden among these poorly studied groups.

Large population sizes can theoretically be sufficient to explain effective purifying selection in asexual oribatid mites, but additional non-mutually exclusive mechanisms could contribute to the reduction of the mutational load. Substitution rates can be altered via molecular mechanisms. For example, gene conversion has prevented mutation accumulation in the human Y chromosome and higher plant chloroplasts ^58,59^ and DNA-repair contributes to the maintenance of DNA integrity in animal mitochondria ^60^. However, as the CDC mainly reflects the impact of purifying selection rather than differences in mutation rates, large population size is likely the most important factor to prevent the accumulation of deleterious mutations in asexual oribatid mites. Finally, some non-canonical form of sex (like e. g. horizontal gene transfer between bdelloid rotifers ^27^) in oribatid mites cannot formally be excluded, albeit it seems unlikely that this would result in more effective selection than common bisexual reproduction.

In conclusion, we conducted several detailed analyses of mutation accumulation in transcriptomes of three asexual and sexual oribatid mite lineages and found purifying selection to be more effective in asexual lineages. This contrasts empirical evidence from younger asexual lineages and non-recombining genome portions. Additional studies in other ancient asexual animal groups would allow testing whether escaping deleterious mutation accumulation is a general feature of ancient asexuals. To what extent population sizes and/or molecular mechanisms contribute to the escape of mutational meltdown remains to be investigated. The results indicate that deleterious mutation accumulation is unlikely to drive extant asexual oribatid mite lineages to extinction and challenge the established consensus that loss of sex necessarily leads to reduced effectiveness of purifying selection and mutational meltdown in the long-term.

## Methods

### Animal sampling, RNA extraction and sequencing of transcriptomes

For analyses of transcriptomic data, we selected three sexual species (*Achipteria coleoptrata:* Brachypylina, *Hermannia gibba:* Desmonomata, *Steganacarus magnus:* Mixonomata) and three asexual species (*Hypochthonius rufulus:* Enarthronota, *Nothrus palustris:* Desmonomata, *Platynothrus peltifer:* Desmonomata) representing four of the six oribatid mite major groups ^34^ (Fig. 1). Animals were collected from litter in different forests in Germany (Göttinger Wald, Solling). Animals were extracted from litter alive using heat gradient extraction ^61^, identified following Weigmann 2006 ^62^ and starved for at least one week to avoid contamination through gut content RNA. For RNA extraction, between 1 and 50 individuals were pooled, depending on body size of the species, to obtain sufficient RNA for library construction. RNA extraction was done using Qiagen and ZymoResearch RNA extraction kits, following manufacturer's instructions and RNA concentrations were measured using the Qubit 2.0 fluorometer (Invitrogen). Library preparation and sequencing of A. *coleoptrata, H. rufulus, N. palustris* and *H. gibba* was performed by GATC Biotech (Constance, Germany); *P. peltifer* and S. *magnus* by the Transcriptome and Genome Analysis Laboratory (TAL, Georg-August-University Göttingen, Germany). Sequencing was done on Illumina platforms yielding 125 or 250 bp paired-end reads. For detailed information, see Supplementary Table 2.

### Transcriptome assembly, ortholog detection and alignment processing

Quality of raw reads was controlled, checked for adapters and the first 12 bases of each read were clipped and bases with quality scores below 30 were trimmed from the end of the reads using Trimmomatic version 0.33 ^63^. Trimmed read pairs were assembled with Trinity version 2.0.3 ^64^ on the scientific computing clusters of the GWDG (Georg-August-University Göttingen). Remains of adaptors and sequences matching to non-oribatid mite species in the ncbi database were removed. The cleaned assemblies were deposited under NCBI BioProject PRJNA339058. To avoid spurious contigs from sequencing errors, only contigs with RPKM > 2 after mapping with RSEM version 1.2.21 ^65^ were selected and the most abundant isoform per gene extracted following Harrison et al. 2015 ^66^. Open reading frames were detected using TransDecoder version 2.0.1 ^67^. For subsequent analyses of branch-specific dN/dS ratios, codon usage bias and hydrophobicity changes, orthologous ORFs were detected using reciprocal best alignment heuristics implemented in Proteinortho version 5.11 ^68^ with amino acid sequences of detected ORFs as input. Amino acid sequences that were shared by all species were aligned with MUSCLE version 3.8.31 ^69^ and back-translated using RevTrans version 1.4 ^70^. Alignments were curated (including deletion of triplet gaps) using Gblocks with parameters −t=c, −b=4 ^71^. Alignments < 200 bp were excluded from further analyses, leaving 3,545 alignments (mean alignment length 943 bp) for dN/dS ratio and codon usage analysis. For detailed information see Supplementary Table 3. Mitochondrial genes were identified with blastn and tblastx and standard parameters ^72^ using unpublished mitochondrial genes derived from genome data and mitochondrial genes of S. *magnus* as query sequences ^73,74^. Alignment and alignment processing was done as described above for nuclear orthologous genes.

For the general phylogeny and the analyses on the increased taxon sampling, we similarly aligned partial coding sequences of *ef1α, hsp82* and 18S rDNA of 30 oribatid mite species, generated in a previous study ^34^. Sequences of these genes were downloaded from NCBI, except for *Nothrus palustris* for which sequences were extracted from the transcriptome data generated in this study using blast (Fig. 1, Supplementary Table 1).

### Tree construction and branch-length estimates for dN/dS ratio analysis

To reconstruct the phylogeny of the large taxon sampling of all 30 species (that includes the six species for which transcriptomes were generated), the alignments of *ef1α, hsp82* and 18S were concatenated and passed to RAxML version 3.1 ^75^ with the GAMMAINVG model set as model of sequence evolution (Fig. 1). For dN/dS ratio analyses, a fixed unrooted species tree, gene-specific alignments and gene-specific branch lengths are required. For nuclear and mitochondrial orthologs derived from the transcriptome data, the 30-taxa topology was compacted to contain the six species only and per-gene branch lengths were calculated using this fixed tree together with the processed alignments with RAxML and the GAMMAINVG model of sequence evolution. Similarly, for the large taxon set, branch lengths of *ef1α* and *hsp82* were calculated using the 30-species tree. For all gene-specific trees, branch lengths were multiplied with three to match the requirements of CodeML for testing models of codon evolution.

### Branch-specific dN/dS ratio analyses

To infer coding mutation accumulation in the three sexual and asexual lineages on transcriptome level, the alignments of 3,545 orthologous genes were analysed using CodeML, implemented in the PAML package version 4.8 ^76^. To get gene-specific dN, dS and dN/dS ratios, a custom script was used to pass each gene alignment together with the fixed species tree, appropriate branch labels (according to the model) and the respective individual branch lengths generated with RAxML to CodeML (available at https://github.com/ptranvan/mites2codeml). CodeML utilizes a Maximum Likelihood framework to estimate the goodness of fit of a codon substitution model to the tree and the sequence alignment and calculates branch-specific dN/dS ratios. First, a model, allowing for one dN/dS ratio for sexual, asexual and internal branches each, was used to calculate dN/dS ratios (three-ratio model). The dN/dS ratios of terminal asexual and sexual branches were compared using a Wilcoxon signed-rank test in R version 3.0.2 ^77^. To investigate if the results are influenced by high non-coding substitution rates rather than by low coding substitution rates, values of dS and dN were compared between terminal sexual and asexual branches using a non-parametric permutation ANOVA with 5000 bootstrap replicates, that does not require specific assumptions for distributions of data (as utilized in previous studies ^15,78^). To identify orthologous genes that are under significantly stronger purifying selection either at asexual or at sexual branches, the goodness of fit of the three-ratio model was compared to that of a model allowing for only one dN/dS ratio at terminal and internal branches, respectively (two-ratio model), using a likelihood ratio test (colored bars, Fig. 2a). Resulting *P* values were false discovery rates adjusted to account for multiple hypothesis testing using the R package qvalue ^79^. To corroborate these analyses with a larger taxon set, branch-specific dN/dS ratios were predicted using the alignments of *ef1α* and *hsp82* with the respective phylogenetic trees. Differences between terminal sexual and asexual branches were tested for significance as described above. To test if high non-coding mutation rates rather than low coding mutation rates affect the results, values of dS and dN were compared between terminal sexual and asexual branches using a t-test after checking for variance similarity between the two groups. Additionally, mean pairwise amino acid divergences were calculated for *ef1α* and *hsp82* to test if divergence was sufficiently high to detect possible differences between dN/dS ratios at terminal sexual and asexual branches.

### Analysis of mutation deleteriousness

To infer the deleteriousness of coding mutations, changes in hydrophobicity at amino acid replacement sites were analysed. To infer the ancestral states of amino acids for the nuclear orthologous genes, first Proteinortho was used to predict orthologous loci among translated ORFs of the six oribatid mite species and the transcriptome of *Tetranychus urticae*, that was used as outgroup ^80^. Orthologous ORFs were aligned using MUSCLE, back-translated using RevTrans and curated using Gblocks with similar parameters as above. Alignments < 200 bp were excluded from further analysis. Branch lengths were calculated for each gene, individually, using the processed alignments with fixed topologies including *Tetranychus urticae* as outgroup and the GAMMAINVG model of sequence evolution. For subsequent prediction of ancestral amino acids, branch lengths of individual trees were multiplied by three and the processed alignments were translated to amino acid using EMBOSS version 6.6.0 ^81^. A custom script was used to pass each translated alignment together with the fixed species tree and individual calculated branch-lengths to CodeML with RateAncestor parameter set to 1 to predict ancestral amino acid sequences for each internal node (https://github.com/ptranvan/mites2codeml). Transitions from ancestral to replacement amino acids were scored for changes in hydrophobicity according to a hydrophobicity scoring matrix ^82^. To infer the ancestral states of amino acids for mitochondrial genes, mitochondrial genes of *T. urticae* were downloaded from NCBI and added to the alignments as outgroup (Accession: NC_010526) ^83^. Alignment and alignment processing was done as described above for nuclear orthologous genes. To infer the ancestral states of amino acids for the large taxon set, sequences of *ef1α* and *hsp82* of the basal oribatid mite *Palaeacarus hystricinus* (Palaeosomata) were downloaded from NCBI and added to the alignments as outgroup. Alignment and alignment processing was done as described above for nuclear orthologous genes. Hydrophobicity transitions were compared between modes of reproduction using generalized LMMs (GLMMs) implemented in the R package lme4 ^84^ with random effect of gene nested in species while correcting for overdispersion and fitted to poisson distribution.

### Analysis of codon usage bias

Bias in codon usage was calculated with the codon deviation coefficient metric (CDC) in Composition Analysis Toolkit version 1.3 ^40^. CDC estimates expected codon usage from observed positional GC and purine contents and calculates the deviation from observed codon usage using a cosine distance matrix, ranging from 0 (no deviation; no detectable (i.e. ‘relaxed’) selection on codon usage) to 1 (maximum deviation; effective selection on codon usage). To analyse codon usage bias of nuclear and mitochondrial orthologous genes, we calculated CDC from processed alignments and compared per gene between reproductive modes using a permutation ANOVA similar to dS and dN comparisons (see above).

### Annotation and enrichment analysis

To analyse enrichment of GO terms in genes that had significantly lower dN/dS ratios either at asexual as compared to sexual branches or at sexual as compared to asexual branches, the 3,545 orthologous genes under purifying selection were annotated for the species *Hypochthonius rufulus* using Blast2GO ^85^. Automatic annotation was successful for 2587 genes, of which 67 genes yielded significantly lower dN/dS ratios at asexual as compared to sexual branches and 5 genes yielded significantly lower dN/dS ratios at sexual as compared to asexual branches.

Based on the 2587 orthologous loci as reference set and the 67 and 5 genes as test sets, enrichment of GO terms was tested for the category Biological Process with Fisher´s exact test in Blast2GO.

### Data availability

The transcriptomic data generated during this study are deposited in NCBI with the BioProject PRJNA339058. Further processed data that support the study are available upon request.

### Code availability

The script for passing the input data to codeml and collecting the dN/dS estimate and hydrophobicity output is available under https://github.com/ptranvan/mites2codeml

## Acknowledgements

We thank Christian Bluhm for help with species identification, Darren J Parker for help with statistics, Patrick Van Tran for help with bioinformatics and Casper van der Kooi, Ken Kraaijeveld, Nicolas Galtier and Roy A. Norton for valuable comments on the manuscript. This study was supported by core funding of S.S., by DFG research fellowship BA 5800/1-1 to J.B. and Swiss SNF grant PP00P3_170627 to T.S.

## Author contributions

A.B. and J.B. conceived and designed the study, A.B., J.G. and J.B. collected samples, A.B. and J.G. performed wet lab work, A.B., J.G. and J.B. performed data analysis, T.S., I.S., M.M., S.S. contributed to data interpretation and analyses and A.B., J.B., I.S. wrote the paper with input from all authors.

## Competing financial interests

The authors declare no competing financial interests.

## Materials & Correspondence

Correspondence to Alexander Brandt and Jens Bast

## References

1. Bell, G. The Masterpiece of Nature: The Evolution and Genetics of Sexuality. 1–635 (Croom Helm Ltd., 1982).

2. Otto, S. P. The evolutionary enigma of sex. Am. Nat. 174, S1–S14 (2009).

3. Williams, G. C. Sex and Evolution. 1-200 (Princeton University Press, 1975).

4. Maynard Smith, J. The Evolution of Sex. 1-222 (Cambridge University Press, 1978).

5. Lehtonen, J., Jennions, M. D. & Kokko, H. The many costs of sex. Trends Ecol. Evol. 27, 172–178 (2012).

6. Jalvingh, K., Bast, J. & Schwander, T. in Encyclopedia of Evolutionary Biology (ed. Kliman R. M.) 89–97 (Academic Press, 2016).

7. Meirmans, S., Meirmans, P. G. & Kirkendall, L. R. The costs of sex: facing real-world complexities. Q. Rev. Biol. 87, 19–40 (2012).

8. Muller, H. J. The relation of recombination to mutational advance. Mutat. Res. 1, 2–9 (1964).

9. Hill, W. G. & Robertson, A. The effect of linkage on limits to artificial selection. Genet. Res. 8, 269–294 (1966).

10. Gordo, I. & Charlesworth, B. The degeneration of asexual haploid populations and the speed of Muller’s ratchet. Genetics 154, 1379–1387 (2000).

11. Lynch, M., Bürger, R., Butcher, D. & Gabriel, W. The mutational meltdown in asexual populations. J. Hered. 84, 339–344 (1993).

12. Johnson, S. G. & Howard, R. S. Contrasting patterns of synonymous and nonsynonymous sequence evolution in asexual and sexual freshwater snail lineages. Evolution 61, 2728–2735 (2007).

13. Neiman, M., Hehman, G., Miller, J. T., Logsdon, J. M. & Taylor, D. R. Accelerated mutation accumulation in asexual lineages of a freshwater snail. Mol. Biol. Evol. 27, 954–963 (2010).

14. Hollister, J. D. et al. Recurrent loss of sex is associated with accumulation of deleterious mutations in Oenothera. Mol. Biol. Evol. 32, 896–905 (2014).

15. Henry, L., Schwander, T. & Crespi, B. J. Deleterious mutation accumulation in asexual Timema stick insects. Mol. Biol. Evol. 29, 401–408 (2012).

16. Hartfield, M. Evolutionary genetic consequences of facultative sex and outcrossing. J. Evol. Biol. 29, 5–22 (2016).

17. Lynch, M. & Blanchard, J. L. Deleterious mutation accumulation in organelle genomes. Genetica 102-103, 29–39 (1998).

18. Bachtrog, D., Hom, E., Wong, K. M., Maside, X. & de Jong, P. Genomic degradation of a young Y chromosome in Drosophila miranda. Genome Biol. 9, R30 (2008).

19. Heethoff, M., Norton, R. A., Scheu, S. & Maraun, M. in Lost Sex - The Evolutionary Biology of Parthenogenesis (eds. Schoen, I., Martens, K. & van Dijk, P.) 241–257 (Springer Netherlands, 2009).

20. Schoen, I., Rossetti, G. & Martens, K. in Lost Sex - The Evolutionary Biology of Parthenogenesis (eds. Schoen, I., Martens, K. & van Dijk, P.) 217–240 (Springer Netherlands, 2009).

21. Mark Welch, D. B., Ricci, C. & Meselson, M. in Lost Sex - The Evolutionary Biology of Parthenogenesis (eds. Schoen, I., Martens, K. & van Dijk, P.) 259–279 (Springer Netherlands, 2009).

22. Judson, O. P. & Normark, B. B. Ancient asexual scandals. Trends Ecol. Evol. 11, 41–46 (1996).

23. Mark Welch, D. B. & Meselson, M. S. Rates of nucleotide substitution in sexual and anciently asexual rotifers. Proc. Natl. Acad. Sci. U. S. A. 98, 6720–6724 (2001).

24. Barraclough, T. G., Fontaneto, D., Ricci, C. & Herniou, E. A. Evidence for inefficient selection against deleterious mutations in cytochrome oxidase I of asexual bdelloid rotifers. Mol. Biol. Evol. 24, 1952–1962 (2007).

25. Debortoli, N. et al. Genetic exchange among bdelloid rotifers is more likely due to horizontal gene transfer than to meiotic sex. Curr. Biol. 26, 723–732 (2016).

26. Signorovitch, A., Hur, J., Gladyshev, E. & Meselson, M. Allele sharing and evidence for sexuality in a mitochondrial clade of bdelloid rotifers. Genetics 200, 1–10 (2015).

27. Schwander, T. Evolution: the end of an ancient asexual scandal. Curr. Biol. 26, R233–R235 (2016).

28. Schoen, I., Martens, K., Van Doninck, K. & Butlin, R. K. Evolution in the slow lane: molecular rates of evolution in sexual and asexual ostracods (Crustacea: Ostracoda). Biol. J. Linn. Soc. Lond. 79, 93–100 (2003).

29. Heethoff, M. et al. High genetic divergences indicate ancient separation of parthenogenetic lineages of the oribatid mite Platynothrus peltifer (Acari, Oribatida). J. Evol. Biol. 20, 392–402 (2007).

30. Von Saltzwedel, H., Maraun, M., Scheu, S. & Schaefer, I. Evidence for frozen-niche variation in a cosmopolitan parthenogenetic soil mite species (Acari, Oribatida). PLoS One 9, e113268 (2014).

31. Maraun, M. et al. Radiation in sexual and parthenogenetic oribatid mites (Oribatida, Acari) as indicated by genetic divergence of closely related species. Exp. Appl. Acarol. 29, 265–277 (2003).

32. Norton, R. A. & Palmer, S. C. in The Acari: Reproduction, Development, and Life-History Strategies (eds. Schuster, R., Murphy, P. W.) 107–136 (Chapman & Hall, 1991).

33. Cianciolo, J. M. & Norton, R. A. The ecological distribution of reproductive mode in oribatid mites, as related to biological complexity. Exp. Appl. Acarol. 40, 1–25 (2006).

34. Domes, K., Norton, R. A., Maraun, M. & Scheu, S. Reevolution of sexuality breaks Dollo’s law. Proc. Natl. Acad. Sci. U. S. A. 104, 7139–7144 (2007).

35. Hershberg, R. & Petrov, D. A. Selection on codon bias. Annu. Rev. Genet. 42, 287–299 (2008).

36. Schaefer, I., Norton, R. A., Scheu, S. & Maraun, M. Arthropod colonization of land - linking molecules and fossils in oribatid mites (Acari, Oribatida). Mol. Phylogenet. Evol. 57, 113–121 (2010).

37. Li, W. H., Wu, C. I. & Luo, C. C. A new method for estimating synonymous and nonsynonymous rates of nucleotide substitution considering the relative likelihood of nucleotide and codon changes. Mol. Biol. Evol. 2, 150–174 (1985).

38. Goldberg, E. E. & Igić, B. On phylogenetic tests of irreversible evolution. Evolution 62, 2727–2741 (2008).

39. Pace, C. N. et al. Contribution of hydrophobic interactions to protein stability. J. Mol. Biol. 408, 514–528 (2011).

40. Zhang, Z. et al. Codon Deviation Coefficient: a novel measure for estimating codon usage bias and its statistical significance. BMC Bioinformatics 13:43, (2012).

41. Ticher, A., Aharon, T. & Dan, G. Nucleic acid composition, codon usage, and the rate of synonymous substitution in protein-coding genes. J. Mol. Evol. 28, 286–298 (1989).

42. Ollivier, M. et al. Comparison of gene repertoires and patterns of evolutionary rates in eight aphid species that differ by reproductive mode. Genome Biol. Evol. 4, 155–167 (2012).

43. Ament-Velásquez, S. L. et al. Population genomics of sexual and asexual lineages in fissiparous ribbon worms (Lineus, Nemertea): hybridization, polyploidy and the Meselson effect. Mol. Ecol. 25, 3356–3369 (2016).

44. Jarne, P. & Auld, J. R. Animals mix it up too: the distribution of self-fertilization among hermaphroditic animals. Evolution 60, 1816–1824 (2006).

45. Goodwillie, C., Kalisz, S. & Eckert, C. G. The evolutionary enigma of mixed mating systems in plants: occurrence, theoretical explanations, and empirical evidence. Annu. Rev. Ecol. Evol. Syst. 36, 47–79 (2005).

46. Maraun, M., Norton, R. A., Ehnes, R. B., Scheu, S. & Erdmann, G. Positive correlation between density and parthenogenetic reproduction in oribatid mites (Acari) supports the structured resource theory of sexual reproduction. Evol. Ecol. Res. 14, 311–323 (2012).

47. Van Doninck, K., Schoen, I., Martens, K. & Goddeeris, B. The life-cycle of the asexual ostracod Darwinula stevensoni (Brady & Robertson 1870) (Crustacea, Ostracoda) in a temperate pond. Hydrobiologia 500, 331-340 (2003).

48. Sahu, G., Gautam, S. K. & Poddar, A. N. Suitable hosts of root knot nematode attack: an assessment on the basis of morphological size variations and population density under field conditions. International Journal of Phytopathology 4, 87–92 (2015).

49. Gaston, K. J., Blackburn, T. M. & Lawton, J. H. Interspecific abundance-range size relationships: an appraisal of mechanisms. J. Anim. Ecol. 66, 579-601 (1997).

50. White, E. P., Ernest, S. K. M., Kerkhoff, A. J. & Enquist, B. J. Relationships between body size and abundance in ecology. Trends Ecol. Evol. 22, 323–330 (2007).

51. Felsenstein, J. The evolutionary advantage of recombination. Genetics 78, 737–756 (1974).

52. Ross, L., Hardy, N. B., Okusu, A. & Normark, B. B. Large population size predicts the distribution of asexuality in scale insects. Evolution 67, 196–206 (2013).

53. Normark, B. B. & Johnson, N. A. Niche explosion. Genetica 139, 551–564 (2011).

54. Rice, W. R. & Friberg, U. in Lost Sex - The Evolutionary Biology of Parthenogenesis (eds. Schoen, I., Martens, K. & van Dijk, P.) 75-97 (Springer Netherlands, 2009).

55. Barton, N. H. Linkage and the limits to natural selection. Genetics 140, 821–841 (1995).

56. Barton, N. H. Why sex and recombination? Cold Spring Harb. Symp. Quant. Biol. 74, 187–195 (2009).

57. Veresoglou, S. D., Halley, J. M. & Rillig, M. C. Extinction risk of soil biota. Nat. Commun. 6:8862 (2015).

58. Hughes, J. F. et al. Strict evolutionary conservation followed rapid gene loss on human and rhesus Y chromosomes. Nature 483, 82–87 (2012).

59. Khakhlova, O. & Bock, R. Elimination of deleterious mutations in plastid genomes by gene conversion. Plant J. 46, 85–94 (2006).

60. Kang, D. & Hamasaki, N. Maintenance of mitochondrial DNA integrity: repair and degradation. Curr. Genet. 41, 311–322 (2002).

61. Kempson, D., Llyod, M. & Ghelardi, R. A new extractor for woodland litter. Pedobiologia 3, 1–21 (1963).

62. Weigmann, G. Hornmilben (Oribatida). Die Tierwelt Deutschlands, begründet 1925 von Friedrich Dahl. 1–520 (Goecke & Evers 2006).

63. Bolger, A. M., Lohse, M. & Usadel, B. Trimmomatic: a flexible trimmer for Illumina sequence data. Bioinformatics 30, 2114–2120 (2014).

64. Grabherr, M. G. et al. Full-length transcriptome assembly from RNA-Seq data without a reference genome. Nat. Biotechnol. 29, 644–652 (2011).

65. Li, B. & Dewey, C. N. RSEM: accurate transcript quantification from RNA-Seq data with or without a reference genome. BMC Bioinformatics 12:323 (2011).

66. Harrison, P. W. et al. Sexual selection drives evolution and rapid turnover of male gene expression. Proc. Natl. Acad. Sci. U. S. A. 112, 4393-4398 (2015).

67. Haas, B. J. et al. De novo transcript sequence reconstruction from RNA-seq using the Trinity platform for reference generation and analysis. Nat. Protoc. 8, 1494–1512 (2013).

68. Lechner, M. et al. Proteinortho: detection of (co-)orthologs in large-scale analysis. BMC Bioinformatics 12:124 (2011).

69. Edgar, R. C. MUSCLE: multiple sequence alignment with high accuracy and high throughput. Nucleic Acids Res. 32, 1792–1797 (2004).

70. Wernersson, R. & Pedersen, A. G. RevTrans: multiple alignment of coding DNA from aligned amino acid sequences. Nucleic Acids Res. 31, 3537–3539 (2003).

71. Castresana, J. Selection of conserved blocks from multiple alignments for their use in phylogenetic analysis. Mol. Biol. Evol. 17, 540–552 (2000).

72. Altschul, S. F., Gish, W., Miller, W., Myers, E. W. & Lipman, J. Basic local alignment search tool. J. Mol. Biol. 215, 403–410 (1990).

73. Domes, K., Maraun, M., Scheu, S. & Cameron, S. L. The complete mitochondrial genome of the sexual oribatid mite Steganacarus magnus: genome rearrangements and loss of tRNAs. BMC Genomics 9:532 (2008).

74. Bast, J. et al. No accumulation of transposable elements in asexual arthropods. Mol. Biol. Evol. 33, 697–706 (2015).

75. Stamatakis, A. RAxML version 8: a tool for phylogenetic analysis and post-analysis of large phylogenies. Bioinformatics 30, 1312–1313 (2014).

76. Yang, Z. PAML 4: Phylogenetic analysis by maximum likelihood. Mol. Biol. Evol. 24, 1586–1591 (2007).

77. R Core Team. R: A Language and Environment for Statistical Computing R Foundation For Statistical Computing https://www.R-project.org/ (2015).

78. Manly, B. F. J. Randomization, Bootstrap and Monte Carlo Methods in Biology. 1-480 (Chapman & Hall/CRC, 1997).

79. Storey, J. D. qvalue: Q-value estimation for false discovery rate control. R package version 2.0.0 https://github.com/StoreyLab/qvalue (2015).

80. Grbić, M. et al. The genome of Tetranychus urticae reveals herbivorous pest adaptations. Nature 479, 487–492 (2011).

81. Rice, P. EMBOSS: the european molecular biology open software suite. Trends Genet. 16, 276–277 (2000).

82. Riek, R. P. et al. Evolutionary conservation of both the hydrophilic and hydrophobic nature of transmembrane residues. J. Theor. Biol. 172, 245–258 (1995).

83. Van Leeuwen, T. et al. Mitochondrial heteroplasmy and the evolution of insecticide resistance: non-Mendelian inheritance in action. Proc. Natl. Acad. Sci. U. S. A. 105, 5980–5985 (2008).

84. Bates, D., Maechler, M., Bolker, B. & Walker, S. Package ‘lme4’: Linear Mixed-Effects Models using 'Eigen' and S4. R package version 1.1-12 https://cran.r-project.org/web/packages/lme4/ (2016).

85. Conesa, A. et al. Blast2GO: a universal tool for annotation, visualization and analysis in functional genomics research. Bioinformatics 21, 3674–3676 (2005).

